## Supplementary data for "LINE1-mediated reverse transcription and genomic integration of SARS-CoV-2 mRNA detected in virus-infected but not in viral mRNA-transfected cells"

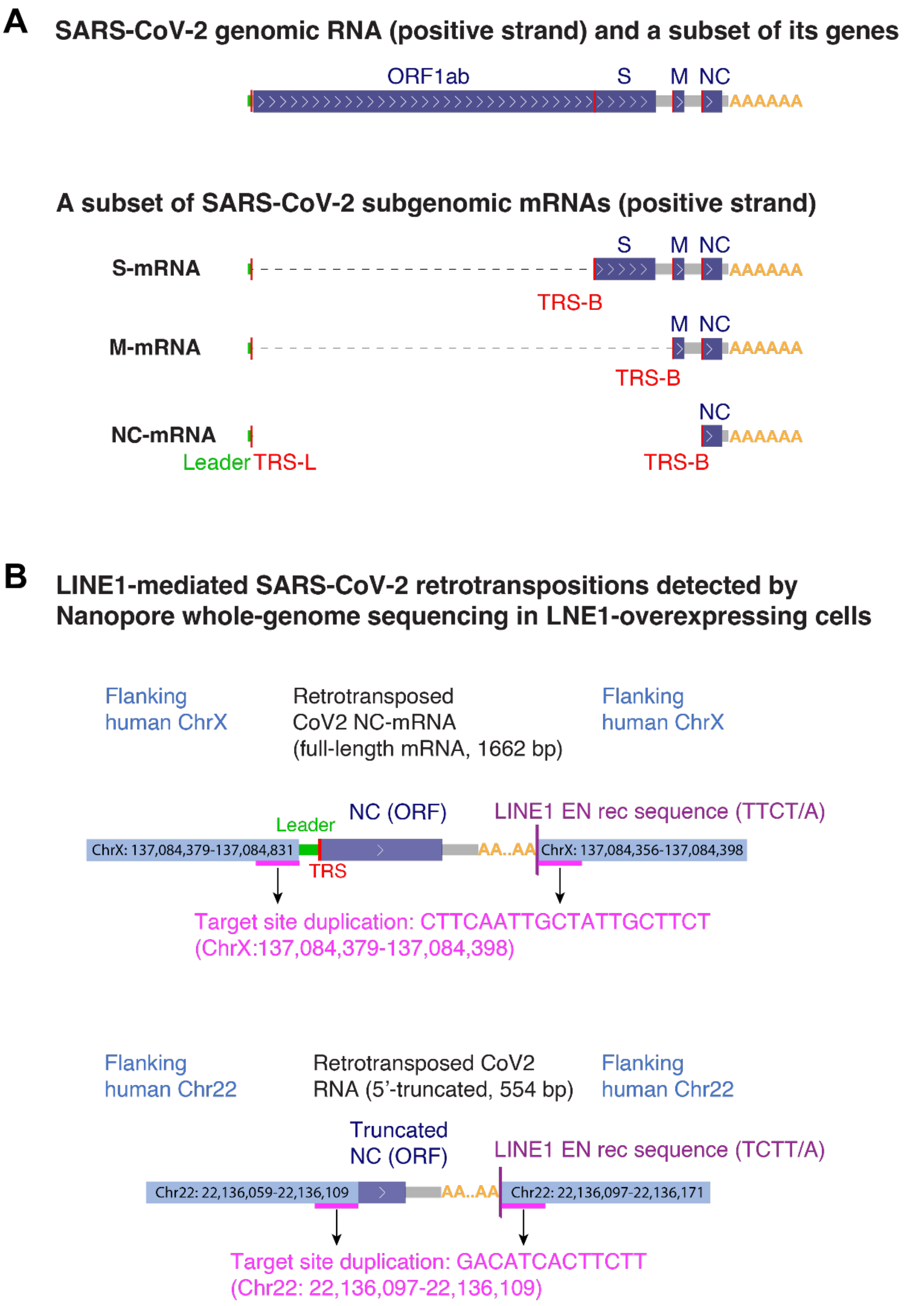


**Supplementary Figure S1. SARS-CoV-2 RNA architecture and retrotransposition by a LINE1-mediated TPRT mechanism. A)** Architectures of SARS-CoV-2 genomic RNA and a subset of subgenomic mRNAs with sequence features of 5'-leader (green), genes (dark blue), transcriptional regulatory sequence leader (TRS-L) or body (TRS-B), and polyadenylation (orange). S: Spike; M: Membrane; NC: Nucleocapsid. **B)** Summary of SARS-CoV-2 RNA retrotranspositions in human chromosome X (top) or chromosome 22 (bottom) with flanking host sequences from both sides recovered by Nanopore WGS, published in reference (27).

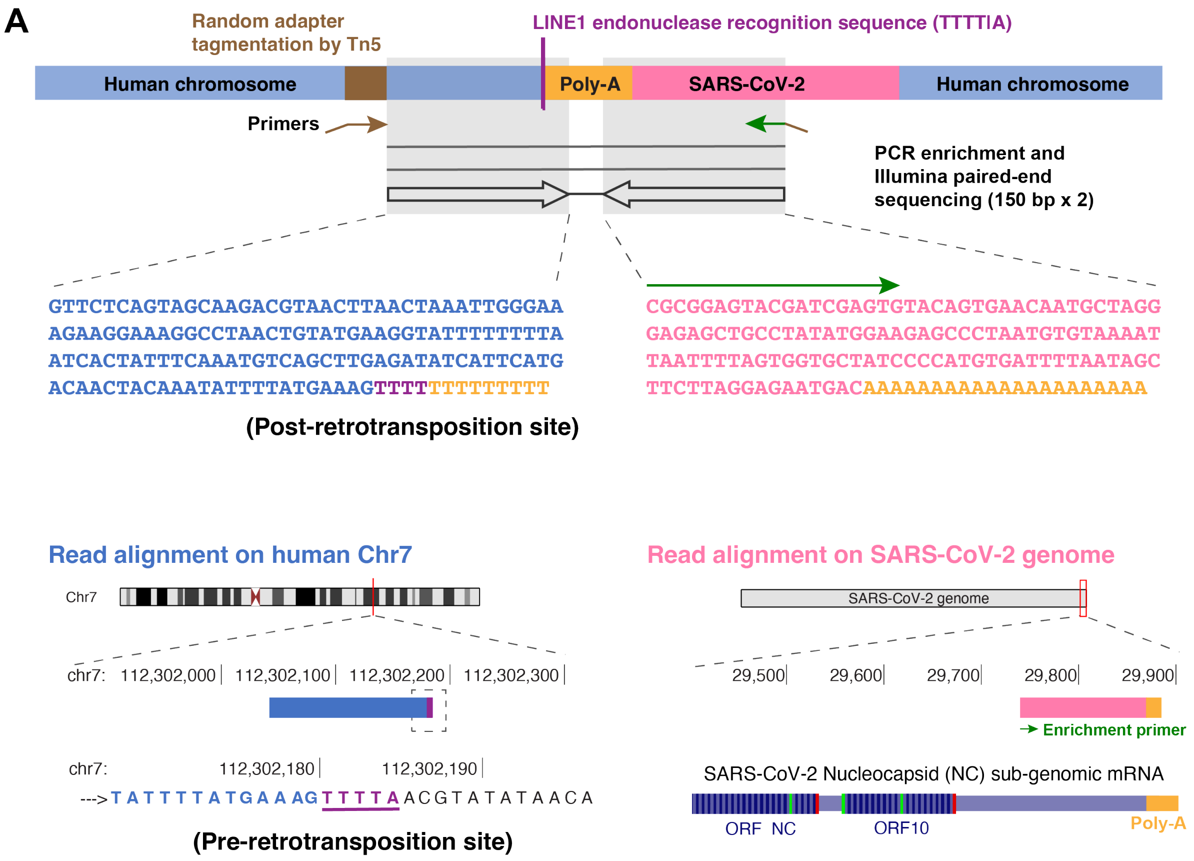


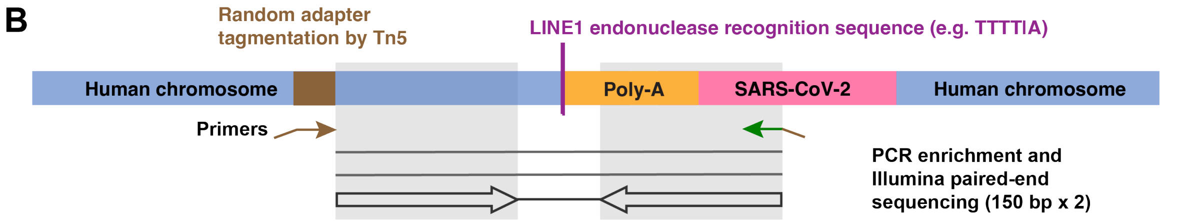


**C**

| Replicate | #1 | #2 |
|---|---|---|
| Integrants with mapped LINE1 EN rec sequence linked to a Poly(A) tract | 33 | 54 |
| Integrants with mapped Poly(A) tract and a putative LINE1 EN rec sequence | 645 | 1056 |
| Total | 678 | 1110 |

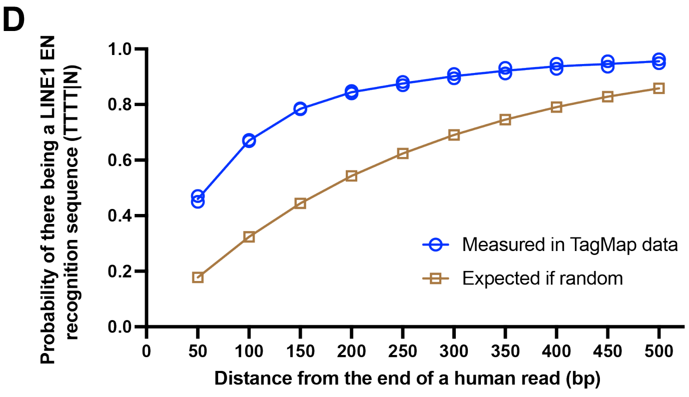

**Supplementary Figure S2. TagMap method for enrichment and sequencing of retrotransposition junctions. A)** Schematic and an example sequencing read-pair showing the enrichment of one junction of retrotransposed viral cDNA. This method is based on random Tn5 tagmentation on cellular genomic DNA. A primer targeting the inserted adapter sequence (brown arrow) and a primer targeting a SARS-CoV-2 sequence (green arrow) are used to enrich the retrotransposition junction. In this example, the left read mapped to human chromosome 7 (blue) with a LINE1 endonuclease recognition sequence (TTTT/A, purple) linked to a poly-A sequence (poly-A tract, orange). The right read mapped to the 3'-end of a SARS-CoV-2 RNA sequence (pink) starting from the enrichment primer sequence (green arrow) and ending with a poly-A sequence derived from the 3'-end viral RNA poly-A (poly-A tract, orange). **B)** Schematic showing the enrichment of one junction of retrotranspositions by a primer (brown) targeting randomly genomic-inserted adaptor and a viral primer (green arrow) targeting the 3'-end of SARS-CoV-2 RNA sequence. In this case, the left read mapped to a human chromosome. The right read mapped to the 3'-end of a SARS-CoV-2 RNA sequence starting from the enrichment primer sequence (green arrow) and ending with a poly-A tract (orange). LINE1 endonuclease recognition sequence is not directly mapped in the read pair due to read length limitation. **C)** Summary of SARS-CoV-2 RNA retrotranspositions detected by TagMap in LINE1-overexpressing 293T cells. All detected retrotranspositions are evidenced by Illumina read-pairs mapped to human sequences and viral sequences with poly-A tract, with or without a LINE1 endonuclease recognition sequence mapped in the reads. **D)** Probability of seeing a LINE1 recognition sequence motif (TTTT) from the mapped human sequence in certain distance intervals (bp), showing measurements in the TagMap data and the expected probability if this motif (TTTT) appears randomly in a given DNA sequence.

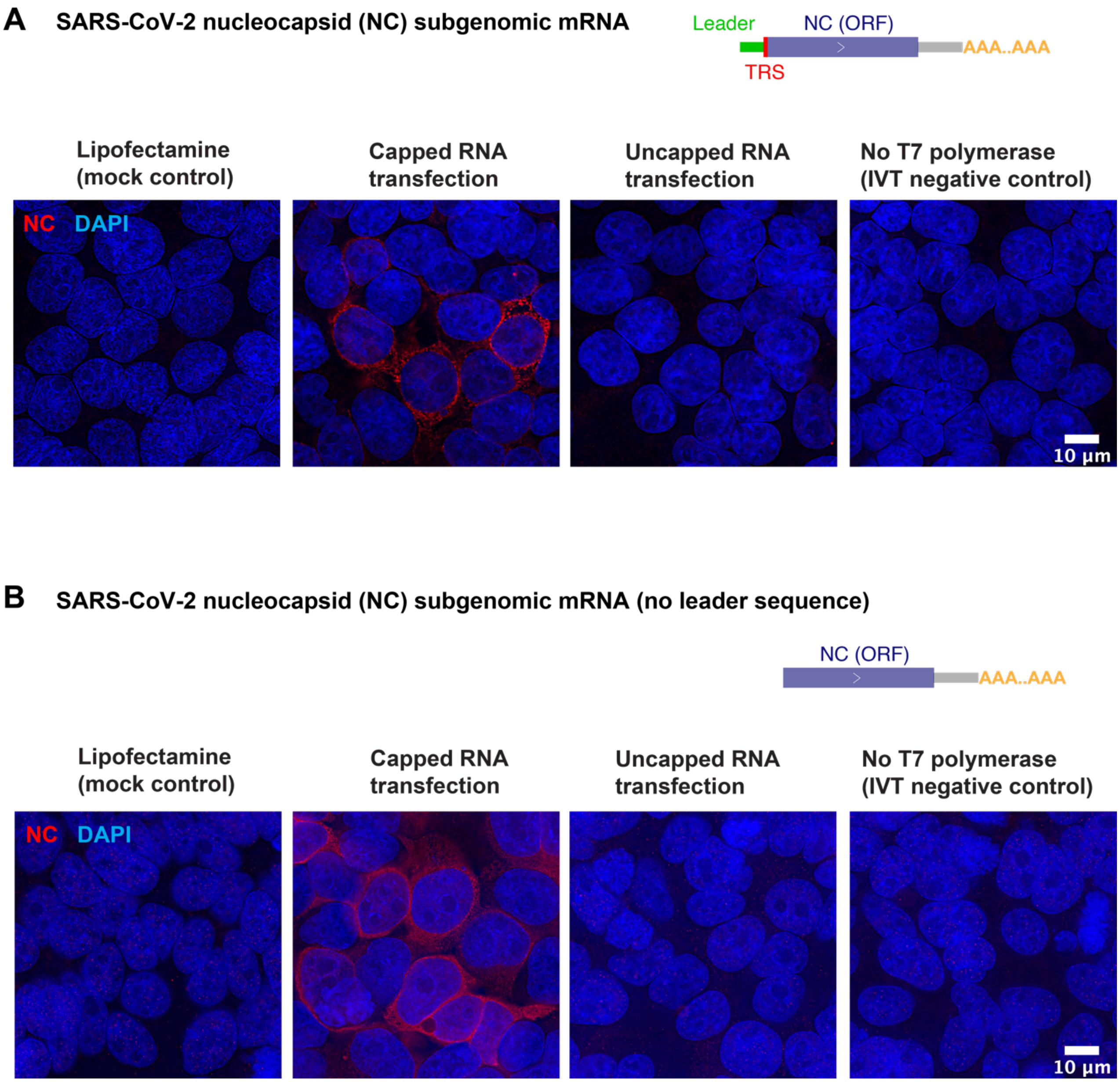


**Supplementary Figure S3. SARS-CoV-2 nucleocapsid mRNA expression in transfected cells. A)** Immunofluorescent staining of nucleocapsid (red) and DNA DAPI staining (blue) in 293T cells transfected by mock, SARS-CoV-2 nucleocapsid mRNA with 5'-cap, SARS-CoV-2 nucleocapsid mRNA without 5'-cap, or an *in vitro* transcription negative control eluent (no T7 polymerase). **B)** Immunofluorescent staining of nucleocapsid (red) and DNA DAPI staining (blue) in 293T cells transfected by mock, SARS-CoV-2 nucleocapsid mRNA (no leader sequence, with 5'-cap), SARS-CoV-2 nucleocapsid mRNA (no leader sequence, without 5'-cap), or an *in vitro* transcription negative control eluent (no T7 polymerase). Cells were transfected by RNA (0.5µg RNA per 1mL cell culture medium) for 24 hours. For IVT negative control, a volume of eluent equivalent to the RNA samples was used for transfection.

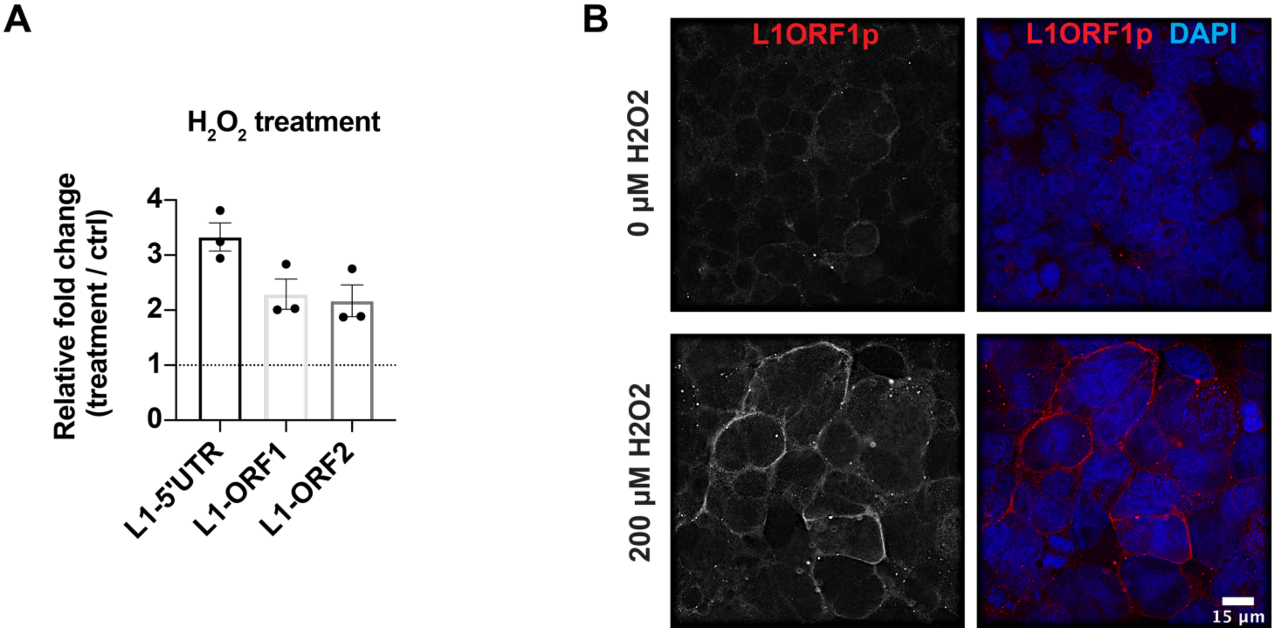


**Supplementary Figure S4. Oxygen stress can induce endogenous LINE1 expression in cultured 293T cells. A)** LINE1 expression fold-change detected by RT-qPCR in 293T cells treated by 200µM hydrogen peroxide for 2 hours followed by 24 hours of culturing. RT-qPCR was performed using L1HS/L1PA(2-6) specific primers targeting LINE1 5'-UTR or ORF1, and L1HS specific primers targeting LINE1 ORF2, on purified cellular poly-A RNA (method and primer sequences following a previous publication (2), see Materials and Methods). n = 3 independent experiments (biological replicates). One-tailed t-test for LINE1 mRNA upregulation in hydrogen peroxide treated cells: p = 0.112 (L1-5'UTR), p = 0.154 (L1-ORF1), p = 0.087 (L1-ORF2). **B)** Immunofluorescent staining of L1ORF1p (red) and merged channels with DAPI staining (blue) in PBS or hydrogen peroxide treated 293T cells. Cells were treated by 200µM hydrogen peroxide for 2 hours followed by 3 days of culturing.
